# The devil lies in the details: small structural and chemical changes in iron oxide pigments largely alter the biological outcomes in macrophages

**DOI:** 10.1101/2025.09.08.674841

**Authors:** Marianne Vitipon, Esther Akingbagbohun, Fabienne Devime, Daphna Fenel, Stéphane Ravanel, Thierry Rabilloud

## Abstract

Because of their technical qualities, such as resistance to fading and to high temperatures, mineral pigments are still widely used nowadays. Among the diversity of mineral pigments, iron oxide pigments represent a widely-used class, because of their technical qualities, diversity of shades (from yellow to red to brown to black) and low toxicity compared to heavy metals-based pigments. However, a low toxicity does not mean the absence of adverse effects. We thus investigated the effects on macrophages of two different subtypes of Pigment Red 101, i.e. hematite, produced by two different processes, namely a wet precipitation process and a calcination process. Macrophages were chosen as a target cell type because they represent the main scavenger cell type that is in charge of handling particulate materials in the body. During this comparison, we realized that the calcined pigment was contaminated from the start by bacterial endotoxins, which induced intense inflammatory responses and biased the comparison. After depyrogenation, the calcined pigment proved to dissolve to a higher extent in macrophages, but to show less important adverse effects (e.g. alteration of the mitochondrial transmembrane potential, oxidative stress and inflammatory responses) than the precipitated pigment in a recovery exposure mode, allowing to investigate delayed effects of the pigments. Thus, despite their identical pigment number, pigments differing in their structure and in their synthesis induce different responses from living cells, even if administrated in equivalent amount. This should be taken into account for some applications, such as tattooing. Moreover, endotoxin contamination should also be checked to increase workers and users safety

## 1. Introduction

Pigments are used in many aspects of our daily life, to bring color to objects. Technically speaking, pigments are insoluble, colored particles that are included into a matrix to produce color on the final object. Nowadays, pigments can be mineral or organic, and organic chemistry has brought a variety of colors that were difficult to reach, at least at a reasonable cost, with mineral pigments only. The technical qualities that are expected from a pigment are mainly its opacity, fluidity and resistance to fading, either chemically- or photo-induced. In some applications, such as plastic coloring or even more severely ceramics, resistance to high temperatures can become a cardinal technical quality. Both in terms of resistance to fading and resistance to heat, mineral pigments offer superior performances, which explains why they are still widely used today.

Before the expansion of the field of organic pigments in the 19^th^ and 20^th^ centuries, mineral pigments were the only source of pigments available. Some of these mineral pigments are present as insoluble mineral deposits, such as cinnabar (HgS), magnetite (Fe3O4), hematite (Fe2O3) azurite or malachite (copper carbonates), or lapis lazuli (complex aluminosilicate containing sulfur). With the development of mineral chemistry over time, many other mineral pigments became available, quite often derived from heavy metals, such as lead white, lead chromate (yellow), cadmium sulfide (from yellow to red), cobalt aluminosilicate (blue).

Although these mineral pigments are endowed with exceptional technical qualities, they come with a major drawback in the form of their toxicity. Pigments are by definition insoluble particles, but only under their intended conditions of use. Acidic body fluids, such as the stomach fluid, easily dissolves carbonates, phosphates and sulfides, although more slowly. This has led to the documented toxicity of lead white (Medley, 1982), which can take surprising turns (Marino *et al*., 1990), or of cinnabar (Davis, 1960; Biro and Klein, 1967; Young *et al*., 2002; Huang *et al*., 2007). Indeed the use of these heavy metals pigments has been associated with health problems in famous painter artists (Pedersen and Permin, 1988).

Nevertheless, there is a family of mineral pigments that are both the most anciently used and considered as the safest mineral pigments, the iron-based pigments. As ochre, which is a mixture of iron oxides, clay and silica, iron oxides are one of the oldest known pigments, and their use traces back to the paleolithic era, although ochre can cover a variety of different pigmentary minerals (Chalmin *et al*., 2021).

Indeed iron pigments can take very different colors, ranging from yellow (goethite, hydrated iron oxohydroxide), to red (hematite, alpha-Fe2O3), brown (maghemite, gamma-Fe2O3) and black (magnetite, Fe3O4). Although ancient time artists only had the raw minerals at their disposal, with the resource of purified mineral salts, they knew that heat treatment could change the color of the iron oxide containing minerals, and especially convert yellow goethite into deep red hematite (Pomiés *et al*., 1999; Cavallo *et al*., 2018). When the art of ceramics developed later, ancient greek artists were able to interconvert hematite into magnetite and vice versa, by changing the oxidizing/reducing conditions in the kiln and by playing with sintering, to produce the famous black-red potteries (Cianchetta *et al*., 2016).

Today, iron oxide pigments are usually prepared from pure salts by wet precipitation routes (Leskelä and Leskelä, 1984; Stefenon *et al*., 2025). However, heat treatment, often called calcination, is still used either to prepare directly hematite, magnetite or maghemite pigments (Leskelä *et al*., 1984; Touzi and Horchani-Naifer, 2023), or to convert pigments prepared by a wet route to heat-stable pigments (Stefenon *et al*., 2025). Thus, pigments designated by the same pigment number (e.g. PR101 for hematite or PBk11 for magnetite) may be produced by either the wet or the thermal processes.

Iron oxide pigments are considered as safe, and thus used in rather invasive applications such as dermopigmentation, the medical form of tattooing (Uhlmann *et al*., 2019; Kuruvilla *et al*., 2022). Although iron oxide pigments do not show the toxicity exhibited by the heavy metal containing pigments (Medley, 1982; Huang *et al*., 2007; Gabby, 1950), this does not mean that they are free of any adverse effects. Indeed, their immediate adverse effects have been documented by several publications on various cell types (e.g. (Luengo *et al*., 2013; Park *et al*., 2014; Dalzon *et al*., 2020; Chrishtop *et al*., 2021)).

Of particular interest are the studies that document their effects on macrophages (e.g. (Park *et al*., 2014; Dalzon *et al*., 2020)), as these cells are in charge of removing all particulate materials from the body, and also play a pivotal role in inflammation. In fact, chronic inflammation driven by persistent particles in macrophages is the driving mechanism for severe pathologies such as asbestosis (Matsuzaki *et al*., 2012) or silicosis (Pollard, 2016). Thus, we recently investigated the delayed effects of iron oxide pigment particles on macrophages, and found adverse effects at the mitochondrial, oxidative stress and inflammation levels (Vitipon, Akingbagbohun, Devime, *et al*., 2025). However, these results were obtained on pigments prepared by a wet route. Thus, we wondered if heat-treated (i.e. burnt or calcined) hematite (PR101) would induce the same delayed effects or different ones on macrophages, as the structure of calcined and un-calcined hematite is known to be different (Pomiés *et al*., 1999).

## 2. Material and methods

### 2.1 Pigment particles preparation and characterization

Pigment Red 101 (red iron oxide, Fe2O3, ref. PS-MI0050, PR101) was purchased from Kama Pigments (Montreal, Canada). Burnt Pigment Red 101 (red iron oxide, Fe2O3, ref. 1675414, PR101B) was purchased from Kremer Pigment (Aichstetten, Germany). When needed, pigments powders were depyrogenized by dry heat treatment in an oven at 220°C for 2 hours (Tsuji and Harrison, 1978), and the pigment designated as PR101BD.

The pigments were received as dry powders and dispersed at 100 mg/ml in an aqueous solution of arabic gum (100 mg/ml), previously sterilised overnight at 80°C in humid atmosphere. Pigment’s dispersions were then re-sterilised under the same conditions to minimise microbial contamination. To reduce agglomerates and aggregates, the dispersions were sonicated in a Vibra Cell VC 750 sonicator (VWR, Fontenay-sous-Bois, France) equipped with a Cup Horn probe. Sonication was carried out in pulse mode for 30 minutes (1 s ON/1 s OFF) at 60% amplitude, corresponding to 90W per pulse (i.e. 81 kJ total energy), in volume not exceeding 3 ml to ensure efficient energy transfer. Dispersions were stored at 4 °C and gently stirred before each experiment to avoid sediment accumulation on tube walls. Prior to use, pigment dispersions were sonicated for 15 minutes in an ultrasonic bath, then diluted in sterile water to intermediate concentrations as required. To limit potential sedimentation and microbial contamination over time, fresh pigment preparations were renewed at least once per month.

For the in-solution characterization of the pigments dispersions, Dynamic Light Scattering (DLS) and Electrophoretic Light Scattering (ELS) were performed using a Litesizer 200 instrument (Anton Paar, Les Ulis, France) equipped with an Omega reusable cuvette (225288, Anton Paar), suitable for both measurements. Before introducing the sample, the zeta potential of the cuvette filled with PBS 0.001X alone was measured to verify cleanliness and exclude background signal or instrument drift. Pigment dispersions were diluted to a final concentration of 50 µg/ml in 0.001X PBS to ensure adequate particle presence for detection while maintaining optimal optical transmittance and ionic strength for electrophoretic mobility analysis. The hydrodynamic size distribution was expressed as intensity-weighted frequency. ELS measurements were conducted at 25 °C, three measurements were taken, each separated by a 30-second delay. Optical parameters were automatically adjusted by the instrument to ensure accurate signal acquisition for each sample.

Transmission Electronic Microscopy (TEM) was performed as previously described (Vitipon, Akingbagbohun, Devime, *et al*., 2025). Briefly, 10 µL of a 100µg/ml dispersion were added to a glow discharge grid coated with a carbon supporting film for 5 minutes. The excess solution was soaked off using a filter paper and the grid was air-dried. The images were taken under low dose conditions (<10 e-/Å2) with defocus values comprised between 1.2 and 2.5 mm on a Tecnai 12 LaB6 electron microscope at 120 kV accelerating voltage using a 4k x 4k CEMOS TVIPS F416 camera.

Finally, the endotoxin concentration of the pigments dispersions was measured with a ToxiSensor kit (GenScript) according to the manufacturer’s instructions.

### 2.2. Cell culture and treatment

The mouse macrophage cell line J774A.1 was purchased from the European Cell Culture Collection (Salisbury, UK). For cell maintenance, cells were cultured in DMEM supplemented with 10% of fetal bovine serum (FBS) in non-adherent flasks (Cellstar flasks for suspension culture, Greiner Bio One, Les Ulis, France). Cells were split every 2 days at 200,000 cells/ml and harvested at 1 million cells/ml.

#### 2.2.1. Determination of Sublethal Dose (LD20)

To determine the sublethal dose 20 (LD20) for each pigment, cells were exposed to a range of pigment concentrations, and viability was assessed using the VVBlue assay (Vitipon, Akingbagbohun, and Rabilloud, 2025) after a 24-hour exposure period. Briefly, J774A.1 macrophages were seeded at 500,000 cells/ml in 24-well adherent plates in DMEM supplemented with 1% horse serum, 1% HEPES, and 10,000 units/ml penicillin-streptomycin as described before (Dalzon *et al*., 2021) to prevent cell proliferation and pigment dilution through time and let rest for 24 hours before pigment exposure. To account for potential interference from pigment absorbance, blank controls containing pigments without cells were included for each condition. This allowed us to correct for background signal in VVBlue assay. Absorbance readings were obtained using a BMG Labtech FLUOstar Omega® plate reader. The LD20 value was defined as the concentration inducing approximately 20% reduction in viability compared to the untreated control.

#### 2.2.2. Recovery exposure settings

Cells were seeded at 500,000 cells/ml in DMEM supplemented with horse serum, as described above. Pigments were added at the selected concentration for 24 further hours as for an acute exposure. Afterwards, the medium was changed in order to remove remaining pigments. The medium was then renewed every 24 hours until the end of the 5 days-recovery period. At the end of the exposure, cells were treated and harvested depending on the functional test performed. For the viability test, the VVBlue assay was used as described above.

### 2.3. Cellular quantification of metals by inductively coupled plasma-mass spectrometry (ICP-MS)

Cells were seeded as previously described into 12-well adherent plates. After the recovery period, cells were rinsed with PBS and harvested into 2 ml Eppendorf tubes. Samples were centrifuged for 5 minutes at 1200 rpm, the supernatant was carefully removed, and 200 µL of ICP lysis buffer (5mM Hepes NaOH pH 7.5, 0.75 mM spermine tetrahydrohloride, 0.1% (w/v) dodecyltrimethylammonio-propane-sulfonate (SB 3-12)) was added to the pellet. The cell pellet was thoroughly vortexed to ensure complete lysis. To separate the soluble and insoluble fractions, lysates were centrifuged at 15,000 g for 30 minutes. The supernatant was transferred into a fresh Eppendorf tube. Samples were stored at -20°C until analysis. The resulting supernatant (soluble fraction) was expected to contain soluble metal ions or degradation products, while the pellet (solid fraction) retained any intracellular, undissolved pigment particles. Both fractions, along with raw pigment suspensions at the working dilution, were analysed by Inductively Coupled Plasma Mass Spectrometry (ICP-MS) to quantify the elemental metal content. Protein concentration in each cell sample was determined using the Bradford assay to normalize metal content to total protein levels.

For the ICP-MS analysis, samples were dehydrated to dryness and digested at 90°C for 4 hours using aqua regia made of three parts of 30% (w/v) hydrochloric acid and one part of 65% (w/v) nitric acid. Mineralized samples were analyzed using an iCAP RQ quadrupole mass spectrometer (Thermo Fisher Scientific). Concentrations were determined using standard curves made from serial dilutions of a multi-element solution (ICP multi-element standard IV, Merck) and corrected using an internal standard solution containing 45Sc and 103Rh, added online. Iron was determined using 56Fe and 57Fe data collected in the helium collision mode. Data integration was done using the Qtegra software (Thermo Fisher Scientific).

### 2.4. Assessment of Cellular Functional Parameters

To evaluate mitochondrial membrane potential and reactive oxygen species (ROS) production, cells were seeded in 12-well adherent plates and exposed to iron-based pigments as previously described. After the recovery period, cells were incubated with their respective probes.

For the mitochondrial membrane potential assay, Rhodamine 123 (Rh123) was added at a final concentration of 10 µM and incubated for 30 minutes at 37 °C. Butanedione monoxime (BDM, 30 mM) was used as a positive control and carbonyl cyanide 4-(trifluoromethoxy)phenylhydrazone (FCCP, 5 µM) as a negative control, both added concurrently with Rh123.

For ROS detection, Dihydrorhodamine 123 (DHR123) was used at a final concentration of 100 µM, with cells incubated for 30 minutes at 37 °C. Menadione was used as a positive control at 50 and 75 µM, added 2 hours prior to probe incubation.

Following incubation with each probe, cold 1X PBS was added to each well, and the plates were incubated for 5 minutes at 4 °C to halt metabolic activity. Approximately two-thirds of the supernatant was discarded after a 3 minutes centrifugation of the plates at 200 g, and the remaining medium was transferred into 2 ml microtubes, one tube per tissue culture well. To collect non-adherent cells, wells were rinsed once with cold PBS and this rinse was added to the corresponding tubes. The tubes were centrifuged at 300 × g for 5 minutes at 4 °C, and the supernatants removed.

In parallel, following plate centrifugation, 200 µL of freshly prepared CTAC lysis buffer (4M urea, 2.5% CTAC, 100mM Hepes NaOH, pH 7.5) was added directly into each well, containing the adherent cells. The plates were placed on ice and gently agitated on a rotating table to ensure complete cell lysis. The lysates from each well were then transferred into the corresponding tubes to lyse the collected non-adherent cells. To separate the soluble protein fraction, samples were then centrifuged at 15,000 × g for 30 minutes at 4 °C. The resulting supernatants were transferred into fresh tubes. For each replicate (N = 4 per condition), 50 µL of the lysate was transferred into a black 96-well plate for immediate fluorescence reading using a BMG Labtech FLUOstar Omega® plate reader (excitation/emission filters: MCB [355-20/460 nm], Rho123 and DHR123 [485-12/520 nm]).). The remaining lysates were stored at −20 °C for subsequent protein quantification using the Bradford assay.

For each assay, corresponding no-probe controls were prepared under identical conditions (same incubation times, treatments, and processing) to evaluate autofluorescence originating from cells, residual pigments, and the lysis buffer. Fluorescence values, with or without probes, were normalized to total protein content. Afterward, without probes controls fluorescence values were subtracted from the corresponding samples to correct for background signals.

### 2.5. Measurement of Cytokines Secretion

To evaluate inflammatory effects either immediately after exposure or following the recovery period, supernatants were transfered into a 2ml tube and centrifuged 5 min at 300g. Afterwards, supernatants were transferred into fresh tubes and Tumor Necrosis Factor alpha (TNFα) and Interleukin-6 (IL-6) levels were quantified using murine ELISA kits (Covalab), according to the manufacturer’s instructions; Absorbances were read using a BMG Labtech FLUOstar Omega® plate reader. The remaining adherent cells were lysed as described above and added to the corresponding tubes containing the non-adherent cells in order to determine total protein content.

In the case of acute exposure, LPS (2 ng/ml) was sometimes added for 24 hours after exposure to the pigments particles, with a medium change.

Cytokines concentrations were normalised to the corresponding protein content of each sample. All experiments were performed in four replicates.

## 3. Results

### 3.1. Physicochemical characterisation of iron-based pigments

The two iron-based pigments in their commercial forms (PR101 and PR101B) together with the depyrogenized pigment (PR101BD) were characterized using Dynamic Light Scattering (DLS) to assess their size distribution, polydispersity index (PI), and sedimentation behaviour.

Thus, the hydrodynamic characteristics of the pigment dispersions were rather similar.

In contrast, representative electron microscopy images in **Figure 1** illustrate the morphological differences between the classical hematite pigment (PR101, Fe2O3; A) and the burnt iron oxide pigment (PR101B, Fe2O3; B). The primary distinction lies in the size of the primary particles. PR101 consists of small spherical particles (<100 nm) that assemble into aggregates, whereas burnt PR101 is composed of larger particles with rather sharp edges, which may be produced by a milling process. Depyrogenation did not alter the shape of the burnt PR101 particles (PR101BD, Figure 1C).

**Figure 1.**
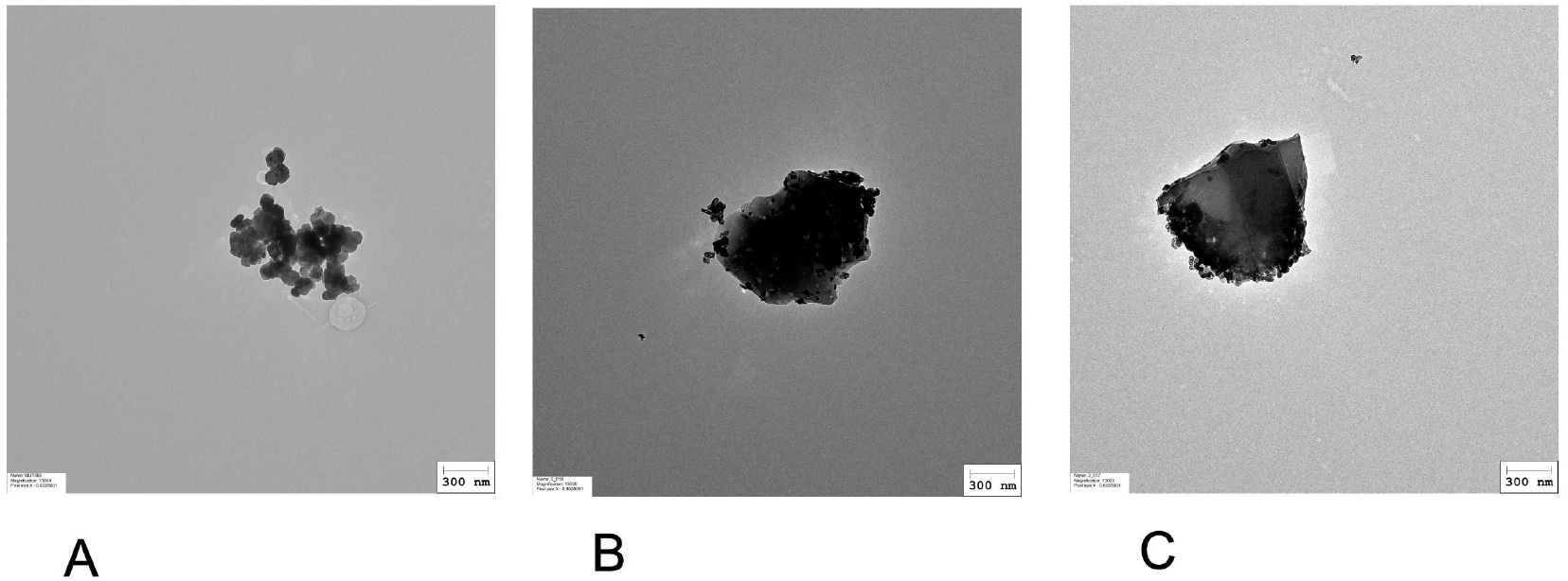
representative electron microscopy images of (A) PR101, (B) burnt PR101 (PR101B), (C) burnt and depyrogenized PR101 (PR101BD). Particles were prepared as described in section 2.1., diluted at 100µg/ml in sterile water. A 10 µL aliquot was applied to a glow discharge grid coated with a carbon supporting film for 5 minutes. The excess solution was soaked off by a filter paper and the grid was air-dried. The images were taken under low dose conditions (<10 e-/Å2) with defocus values between 1.2 and 2.5 µm on a Tecnai 12 LaB6 electron microscope at 120 kV accelerating voltage using 4k x 4k CEMOS TVIPS F416 camera.

### 3.2. Cell exposure and dose selection

For pigment exposure, cells were treated as previously described (Vitipon, Akingbagbohun, Devime, *et al*., 2025).

In a first series of experiments, we determined which concentrations of each pigment could be used on macrophages, by performing viability assays after the acute exposure. In **Figure 2A**, the VVBlue assay (Vitipon, Akingbagbohun, and Rabilloud, 2025) chosen to avoid optical interference from pigment opacity, did not reveal any cytotoxicity of the PR101 pigment for up to 1400 µg/ml. In contrast, as-received burnt PR101 (PR101B) showed some toxicity at 1000µg/ml and above, which disappeared when the pigment was depyrogenized (PR101BD)

**Figure 2.**
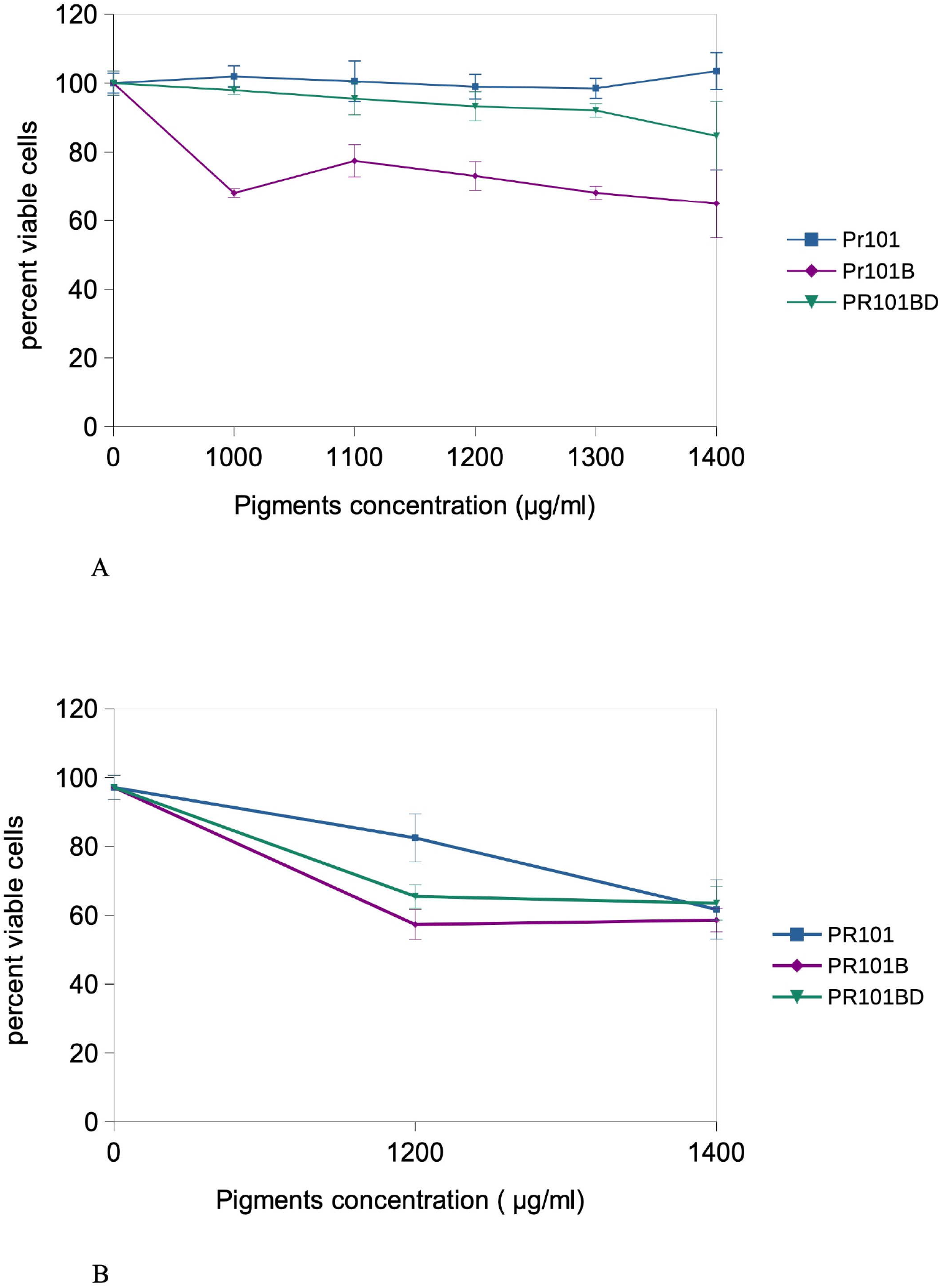
Exposure scheme and viability determination Panel A: viability curve immediately after exposure to pigments (24 hours) blue: PR101 purple: burnt PR101 (PR101B) green: burnt and depyrogenized PR101 (PR101BD) Results are expressed as mean ± standard deviation (N=4) Panel B: viability curve after exposure to pigments for 24 hours followed by 5 days of recovery in a pigment-free medium blue: PR101 purple: burnt PR101 (PR101B) green: burnt and depyrogenized PR101 (PR101BD) Results are expressed as mean ± standard deviation (N=4)

We then tested the two highest concentrations (1200 and 1400 µg/ml) in the recovery format (**Figure 2B**). After 5 days post exposure, PR101 showed a 23% lethality at 1200 µg/ml and a 40% lethality at 1400µg/ml. Burnt PR101 (PR101B) showed a lethality of 40% at both concentrations, while depyrogenized burnt PR101 (PR101BD) showed a lethality of 35% at 1200 µg/ml and a 40% lethality at 1400µg/ml.

Based on these results, 1200 µg/ml was selected for subsequent experiments, which were carried out mostly in the recovery format.

### 3.3. Initial experiments in acute exposure mode

As we knew from previous experiments that PR101 induced a proinflammatory response, we tested the burnt PR101 grade for this response in the acute exposure mode (24 hours exposure to pigments, readout after an additional 24h in a pigment-free medium). The results, shown in **Figure 3**, showed a marked inflammatory response to burnt PR101, even exceeding the one induced by the positive control (LPS (lipopolysaccharide) at 2ng/ml), and with a synergistic effect between burnt PR101 and added LPS (for IL-6). For TNF, the response to burnt PR101 was much less marked. This prompted us to test if the burnt pigment could be contaminated with endotoxins. The endotoxin test yielded a contamination level equivalent to 1.2 pg LPS/g of pigment for PR101, while 90pg/g of pigment were measured for burnt PR101. Depyrogenation of PR101B for 2 h at 220°C reduced the endotoxin level to 12 pg/g of pigment.

**Figure 3.**
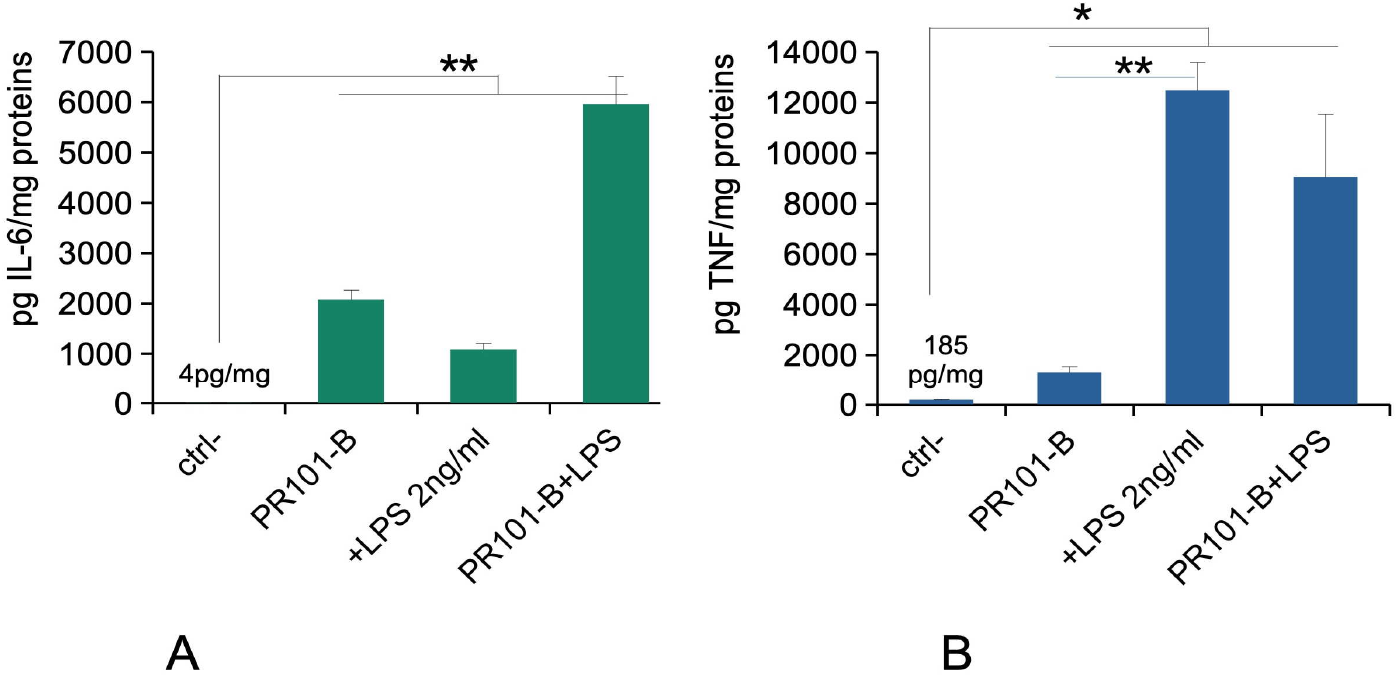
Cytokine release of the raw burnt pigment The cells were exposed to pigments for 24 hours. The cells were treated (or not) with 2 ng/ml lipopolysaccharide in complete, pigment-free cell culture medium for an additional hours. The cell medium was then collected for secreted TNF and IL-6 measurements. Results are expressed as mean±standard deviation (N=3). Significance marks: * = p<0.05; ** = p<0.01 Panel A: IL-6 release under the acute exposure mode Panel B: TNF-alpha release under the acute exposure mode

As this surprising inflammation result could be attributed to the presence of LPS in the pigment, which was greatly reduced by depyrogenation by dry heat, which did not alter the other pigment characteritistics, all subsequent experiments were carried out on the depyrogenized burnt pigment (PR101BD).

### 3.4. Pigments uptake and dissolution by cells

In order to obtain a first appraisal of pigment uptake by macrophages, we performed a simple examination of pigments-loaded cells by optical microscopy. The results, displayed on **Figure 4**, showed that the cells were heavily loaded with PR101 particles, even after the recovery period. This heavy loading rendered the cells opaque, which explains why we could not apply any optical method, and especially flow cytometry, on the pigments-loaded cells.

**Figure 4.**
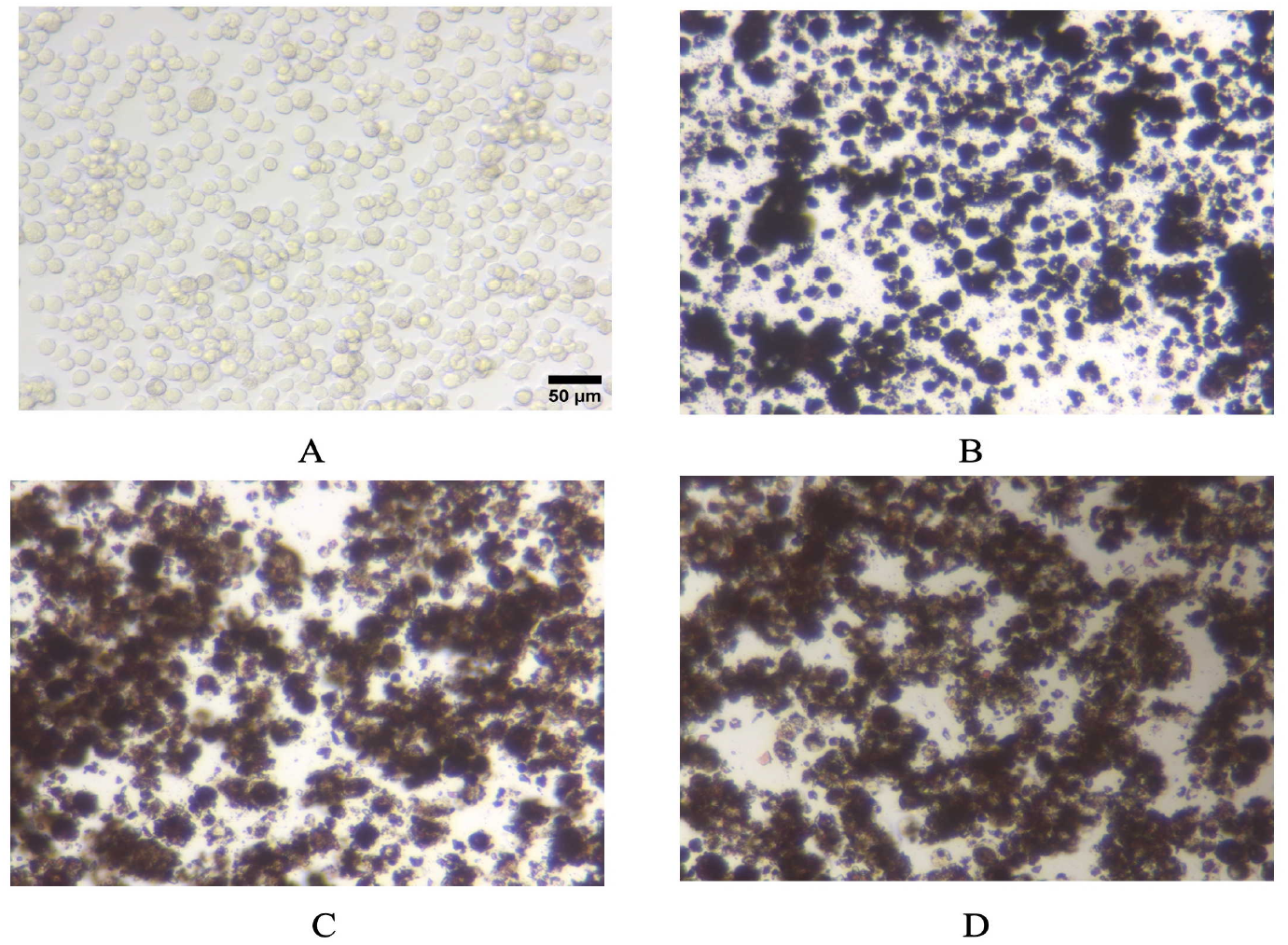
Optical microscopy images of A: control, untreated cells, B: PR101-treated cells C: burnt PR101-treated cells (PR101B) D: burnt and depyrogenized PR101-treated cells (PR101BD)

To more quantitatively assess pigment uptake after acute exposure and retention following the recovery period, mineralization experiments were conducted on the lysates of the cells loaded with PR101BD at 1.2 mg/ml. The supernatant (soluble fraction) gave the proportion of soluble iron, while the pellet (internalized particular fraction) gave the amount of pigment remaining as particular material after the recovery period.

The results, shown in **Table 2**, showed that the burnt and depyrogenized pigment was efficiently internalized by the macrophages, but poorly dissolved even after 5 days of presence in the cells.

**Table 1:** Physicochemical characterization of iron oxide pigments.

|  | **PR101** | **PR101B** | **PR101BD** |
| --- | --- | --- | --- |
| Mean Hydrodynamic diameter (nm) | 386.71 ± 3.44 | 417.06 ± 18.56 | 427.60 ± 22.57 |
| Polydispersity index (%) | 18.79 ± 3.07 | 24.95 ± 0.73 | 26.31 ± 0.92 |
| Zeta potential (mV) | -31.46 ± 0.08 | -30.13 ± 0.22 | -29.81 ± 0.06 |

**Table 2:**
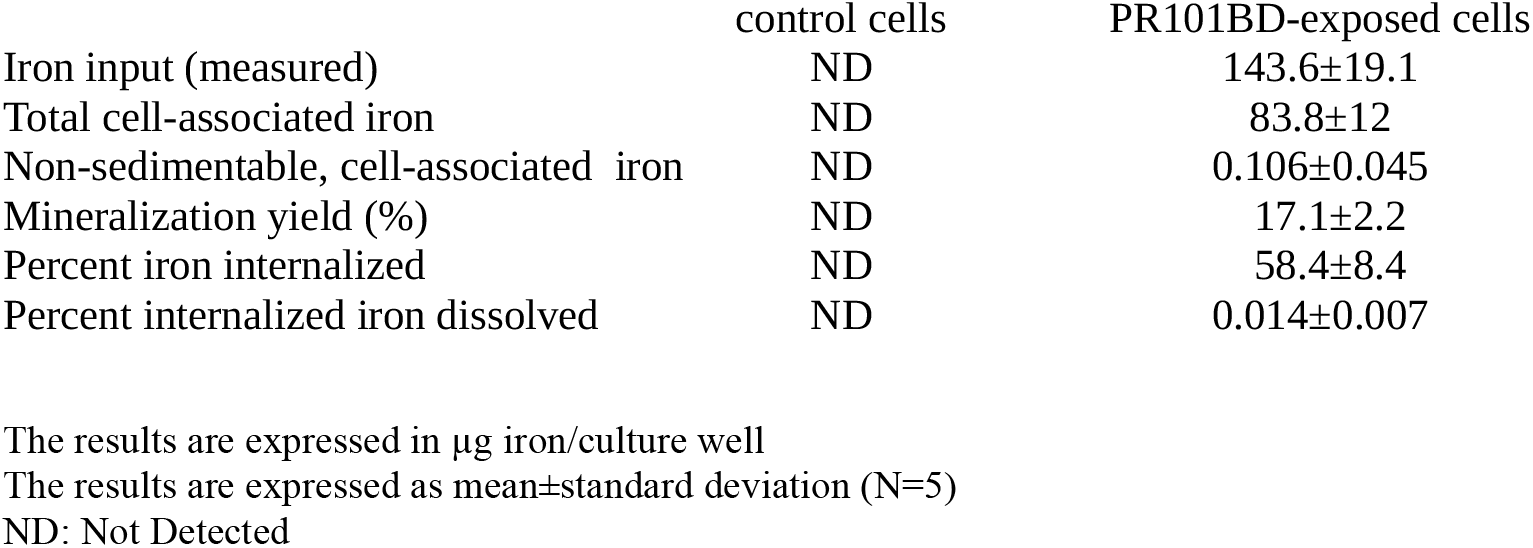
quantification of internalized and dissolved iron in pigments-exposed macrophages after 24 hours of exposure and 5 days of recovery.

In subsequent experiments, the cellular parameters to be tested were selected on the basis of our previous results obtained on PR101 and PBk11 (Vitipon, Akingbagbohun, Devime, *et al*., 2025).

### 3.5. Mitochondria

The mitochondrial transmembrane potential is a key figure of cellular homeostasis. We thus measured it in response to treatment to our various grades of PR101 pigments, in the recovery exposure scheme. The results, displayed in **Figure 5**, showed that PR101 and PR101B (with its adsorbed LPS) significantly increased the mitochondrial transmembrane potential. However, once depyrogenized, the PR101BD pigment did not induce a significant increase of the mitochondrial transmembrane potential.

**Figure 5.**
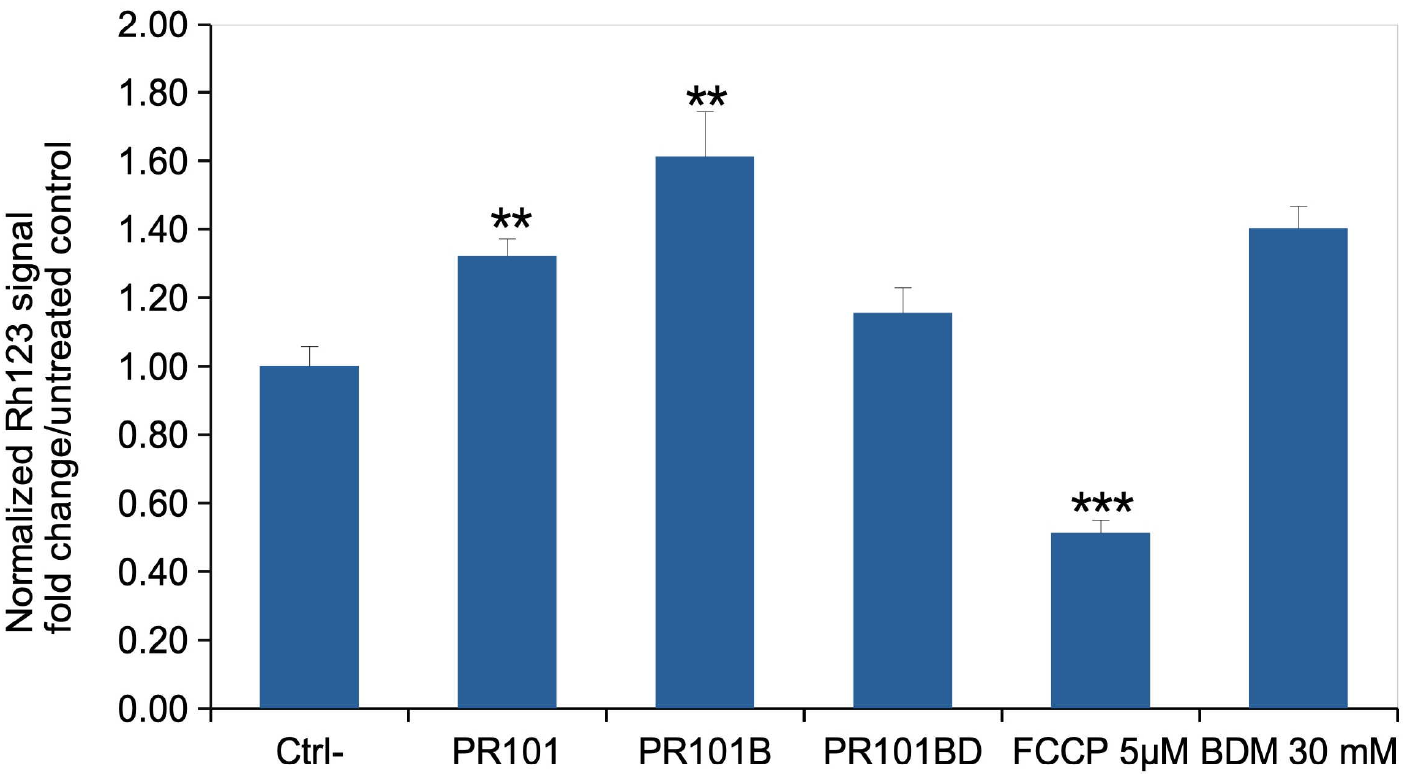
Mitochondrial transmembrane potential The cells were exposed to pigments for 24 hours, then left to recover for 5 days post exposure. Mitochondrial transmembrane potential was then measured via the rhodamine 123 method. All cells were positive for rhodamine 123 internalization in mitochondria, and the mean fluorescence is the displayed parameter. Results are displayed as mean± standard deviation (N=4) CTRL: Unexposed cells PR101: cells exposed to Fe_2_O_3_ pigment PR101B: cells exposed to burnt Fe_2_O_3_ pigment PR101BD: cells exposed to burnt and depyrogenized Fe_2_O_3_ pigment FCCP: Cells exposed for 30 minutes to FCCP (induces a decrease in the transmembrane potential) BDM: Cells exposed for 30 minutes to Butanedione monoxime (induces an increase in the transmembrane potential) Significance marks (pigment vs control): ** = p<0.01; *** = p<0.001

### 3.6. Oxidative stress

As an increase in mitochondrial transmembrane potential has been correlated to oxidative stress (Korshunov *et al*., 1997; Starkov and Fiskum, 2003), we probed the level of oxidative stress in pigment-exposed cells after the recovery phase. The results, displayed in **Figure 6**, showed the same trend than for mitochondrial potential. A moderate but statistically significant increase in response to PR101 and PR101B, disappearing after depyrogenation.

**Figure 6.**
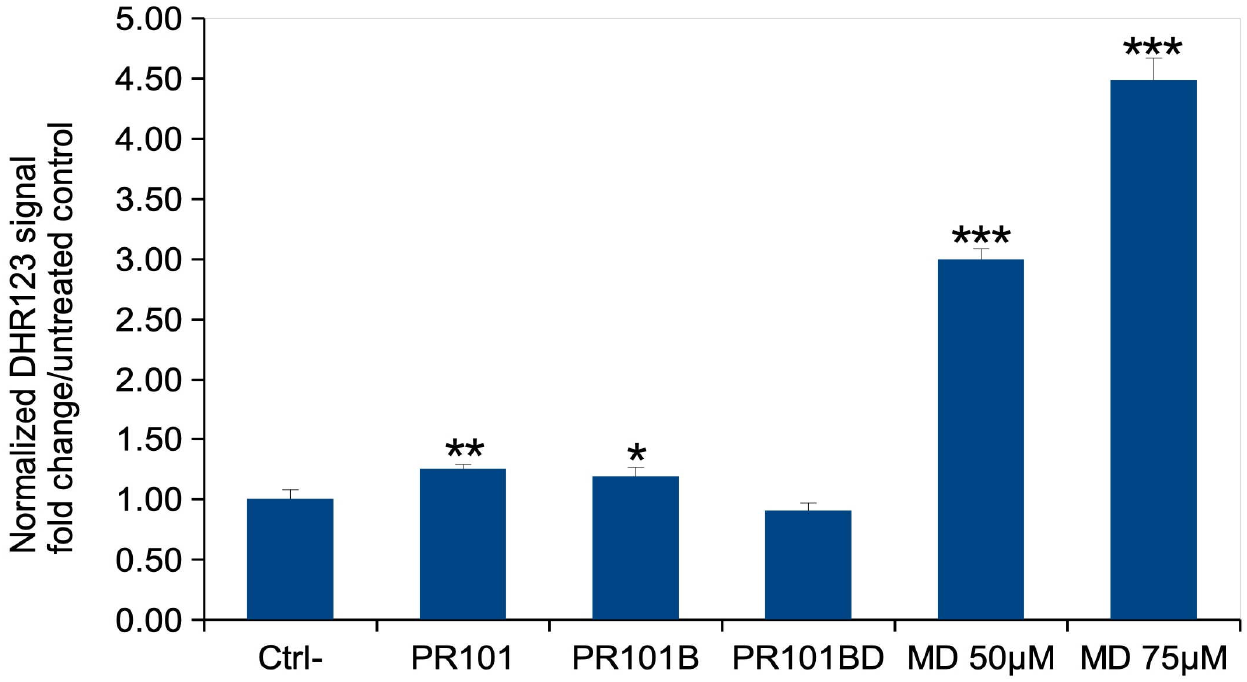
Oxidative stress The cells were exposed to pigments for 24 hours, then left to recover for 5 days post exposure. The level of oxidative stress was measured via the DHR123 method. Results are displayed as mean± standard deviation (N=4). CTRL: Unexposed cells PR101: cells exposed to Fe_2_O_3_ pigment PR101B: cells exposed to burnt Fe_2_O_3_ pigment PR101BD: cells exposed to burnt and depyrogenized Fe_2_O_3_ pigment MD: menadione (added 2 hours before measurement) Significance marks (pigment vs. control): * = p<0.05; ** = p<0.01; *** p<0.001

### 3.7. Immune functions

As our initial findings after acute exposure were puzzling and drove us to explore successfully the LPS contamination hypothesis, we then probed the effect of depyrogenized burnt PR101 on the inflammatory responses of macrophages, in the acute and exposure-recovery formats. The results, displayed in **Figure 7**, showed a general trend in which both PR101 and PR101BD showed an intrinsic pro-inflammatory power, both in the acute exposure and delayed response (post-recovery) format, with PR101BD inducing a lesser inflammatory response than PR101. There was an exception to this tendency in the case of the acute response for IL-6, for which the depyrogenized pigment still induced a higher response than PR101. However, as Il-6 secretion is extremely sensitive to LPS, this might be due to the low residual LPS content even after depyrogenation.

**Figure 7.**
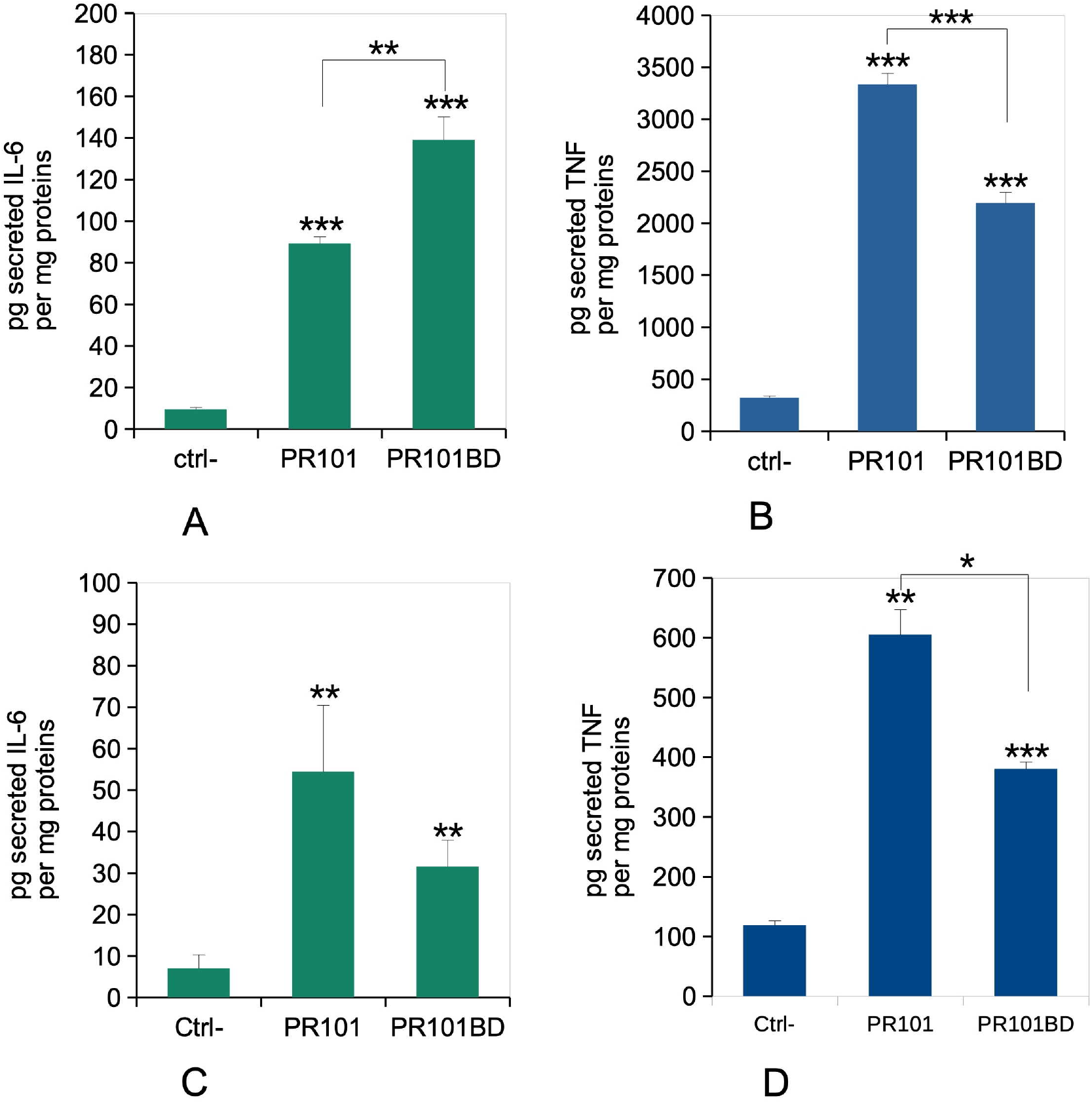
Cytokine release The cells were exposed to pigments for 24 hours, then analysed immediately after exposure (acute mode), or left to recover for 5 days post exposure prior to analysis (recovery mode). The cell medium was then collected for secreted TNF and IL-6 measurements. Results are expressed as mean±standard deviation (N=4). Significance marks (pigment vs. control unless otherwise noted): * = p<0.05; ** = p<0.01; *** p<0.001 Panel A: IL-6 release under the acute exposure mode Panel B: TNF-alpha release under the acute exposure mode Panel C: IL-6 release under the recovery exposure mode Panel D: TNF-alpha release under the recovery exposure mode

## 4. Discussion

The first, unexpected result that we faced in this work was the puzzling strong inflammatory response induced by the burnt iron oxide. Such isolated inflammatory responses have been reported in the past for other nanoparticles such as titanium dioxide (Huang *et al*., 2017). However, the interesting feature of IL-6 is that its secretion is very sensitive to LPS presence, which oriented us to search for contamination of the burnt iron oxide with endotoxins, an approach that was not undertaken in the case of the reported strong pro-inflammatory effects of titanium dioxide (Huang *et al*., 2017). We indeed found a contamination of the pigment by endotoxins, but at low levels (90pg/g). However, it must be kept in mind that the endotoxins are present in the pigment powder, i.e. probably adsorbed at the surface of the pigment particles. We can then only assume that the endotoxin detection kit, based on the limulus amoebocyte lysate, i.e. a coagulation cascade (Young *et al*., 1972), detects the endotoxin similarly to the macrophages, in the sense that the interferences due to endotoxin adsorption on the particles are similar in both cases. If this is the case, then it is striking to observe that these low levels of adsorbed LPS are much efficient than soluble LPS to induce macrophages responses. In the case of IL-6 these ca. 100pg/ml of adsorbed LPS are as efficient as 2 ng/ml of soluble LPS (this work). Regarding toxicity, adsorbed LPS seems much more efficient in inducing J774A.1 cells death than free LPS, which is efficient only in the high ng/ml range (Nikolova *et al*., 2020).

Such LPS-induced inflammatory responses are not a problem in many pigments applications, e.g. in paints when the pigments are embedded in the paint matrix. However, they can become a problem if workers or users inhale such contaminated powders, as it will produce an inflammatory response in the lung through the stimulation of alveolar macrophages. It can also be a problem in the case of usage of such contaminated pigments for tattoo inks, not only at the injection site, where a strong inflammatory reaction is expected in all cases, but also at the draining lymph nodes, where pigments particles are known to accumulate (Schreiver *et al*., 2017), with still debated health consequences (Nielsen *et al*., 2024). In this context, it is surprising to realize that endotoxin levels are not included in regulations on tattoo inks. Sterility is, but sterilization of the inks, e.g. by ionizing radiations, may at the same time succeed in killing all microbes and fail to destroy endotoxins, which are fairly resistant to ionizing radiations and require very high doses to be destroyed (Brydon, 2023). It seems that the regulatory bodies focus on the long term chemical risk of pigments impurities and degradation products, and do not take enough into account the biological risk represented by endotoxins present in inks, maybe as it is perceived as transient in nature. Although it may be well the case for tattoos, it may be different for workers using contaminated pigments powders and thus repeatedly exposed to endotoxins via the contaminated pigments.

It is also interesting to figure out when such bacterial contamination may occur. Burnt iron oxide being a calcined pigment, i.e. fired in an oxidizing atmosphere at several hundred degrees for at least one hour (Leskelä *et al*., 1984; Stefenon *et al*., 2025), endotoxins will be completely destroyed during this step of the pigment synthetic process. This means in turn that contamination can only occur during downstream steps of the process, such as milling or storage. This shows where progress can be made to decrease the endotoxin contamination issues.

Once we reduced the endotoxin level via a subsequent depyrogenation step, we could study the influence of the pigment structure on the macrophages responses. Indeed, iron oxide pigments are usually prepared by wet precipitation reactions and can take several structures and colors (from yellow to black), depending on the synthetic conditions and iron salt reactants (Stefenon *et al*., 2025; Leskelä and Leskelä, 1984). However, these pigments obtained by precipitation can be modified by heat treatments (e.g. (Stefenon *et al*., 2025)). These thermal treatment interconvert pigments between one another, depending on the conditions. Thermal treatments of goethite (yellow iron oxide, PY42) and magnetite (mixed valence iron oxide, PBk11) under an oxidizing atmosphere produce deep red hematite pigment (PR101) (Stefenon *et al*., 2025). Conversely, thermal treatment of goethite (Da Guarda Souza *et al*., 2020) and hematite (Pineau *et al*., 2006; Ponomar, 2018) under reducing atmospheres produces magnetite. Thus, PR101 can be produced with or without thermal treatments. In the case of the pigments tested in this work, thermal treatments drastically altered the morphology of the pigments particles. Instead of the low diameter ovoid or rod like particles obtained by the wet route (this work and, e.g. (Dalzon *et al*., 2020)), burnt PR101 showed more spheric elementary particles, which seem to be produced by the fusion of very small elementary particles, which still can be seen on some edges of the fused particles (see Figure 1). Interestingly, the intracellular dissolution of this burnt and depyrogenized PR101 pigment was intermediate between the low dissolution of unburnt PR101 and the higher intracellular dissolution of magnetite (PBK11) (Vitipon, Akingbagbohun, Devime, *et al*., 2025).

Using this burnt pigment, we compared its effects on macrophages with the most salient effects induced by unburnt PR101, namely mitochondrial dysfunction, oxidative stress and inflammatory reaction (Vitipon, Akingbagbohun, Devime, *et al*., 2025). For all these parameters, we found an inverse correlation between the amount of dissolved iron and the parameter change compared to control. Unburnt PR101, which showed minimal dissolution, showed the highest functional effects, while PBk11, which showed the highest dissolution rate, showed no functional effects on these tested parameters. Burnt and depyrogenized PR101, which showed intermediate dissolution, also showed intermediate functional effects. We thus come to the observation that the adverse effects are inversely proportional to the extent of intracellular dissolution, as if the dissolved iron was triggering protective mechanisms against adverse phenomena induced by the particles themselves.

Interestingly, the endotoxin-contaminated burnt pigment showed higher effects than unburnt PR101 on the mitochondrial potential, but not on oxidative stress.

However, the cellular effects of the endotoxin-contaminated burnt pigment were higher than those of the depyrogenized pigment, which is consistent with the known strong effects of LPS on these parameters on J774A.1 cells (Raza *et al*., 2014).

In conclusion, it appears that pigments bearing the same pigment number are not equivalent, and not only in their technical qualities (e.g. opacity, fluidity, covering power). They are also different in the biological consequences that they induce on the cells that come in contact with them, such as macrophages. The differences in the biological outcome depend on the structure of the pigment particles themselves, but also on spurious phenomena such as contamination of the pigments with exogenous compounds such as endotoxins, as described in this paper. It is thus advisable to check for the presence of potential biologically-effective contaminants, both on the basis of the intended use (e.g. in tattooing) and for the sake of workers safety.

## Acknowledgments

We thank Guy Schoehn for the establishment of the IBS/ISBG EM facility

## Data availability

The cell biology data are available through the BioStudies database under the identifier **S-BSST2099**

## Authors contributions

MV, EA, FD, DF: investigation, formal analysis

SR, TR: formal analysis, funding acquisition, project administration

MV, SR, TR: Writing – original draft

TR: conceptualization

All co-authors: Writing – review and editing

